# Enhanced neural representation of reach target direction for high reward magnitude but not high target probability

**DOI:** 10.1101/2023.12.13.571560

**Authors:** B Keane, E Reuter, J Manzone, B Miller-Mills, L Leow, TN Welsh, TJ Carroll

## Abstract

Many characteristics of goal-directed movements, such as their initiation time, initial direction, and speed, are influenced both by the details of previously executed movements (i.e. action history), and by the degree to which previous movements were rewarded or punished (i.e. reward history). In reinforcement learning terms, when movements are externally cued, action and reward history jointly define the probability and magnitude of positive/negative outcomes of available options, and therefore their pre-stimulus expected value. To dissociate which of these neurocomputational variables influence sensorimotor brain processing, we studied how reach behaviour and evoked brain responses are affected by independent manipulations of action and reward history. We found that movements were initiated earlier both for more frequently repeated targets and targets associated with higher reward magnitude, but only movements to highly rewarded targets had higher movement speeds. Classical visually-evoked encephalographic (EEG) potentials (P1/N1) were not affected by either reward magnitude or target probability. There were, however, amplified midline ERP responses at centroparietal electrodes for rewarded targets and movements compared to control, but no differences between more frequently presented targets and control. Critically, the spatial precision of decoded target locations extracted from a multivariate linear decoding model of EEG data was greater for target locations associated with higher reward magnitude than for control target locations (∼150-300ms after target presentation). Again, there were no differences in the precision of decoded target direction representations between more frequent target locations and control target locations. These data suggest that the expected reward magnitude associated with an action, rather than its long-run expected value, determines the precision of early sensorimotor processing.

**Significance Statement:** We move more quickly and more accurately toward goals that we value more highly, and this is due partly to enhanced motor preparation. However, our expectations about the value of an action depend both on the probability of its requirement and the magnitude of the reward associated with it. Here we disentangled the influence of reward magnitude and probability on early sensorimotor processing via a multivariate linear decoding approach to extract target direction from scalp encephalograms. We found that the spatial precision of decoded target direction was greater for high reward targets but not for more probable targets. Thus, early sensorimotor processing is sharpened when the magnitude of reward associated with movement to a cued target is high.

**Highlights:**

- The direction of movement can be reliably decoded from the scalp EEG from ∼80ms after target presentation.
- The neural representation of movement direction is more precise for targets that are associated with high reward, but not for targets that are more probable.
- The magnitude of reward associated with movement to a presented target, rather than the long-run expected value of the movement, sharpens the spatial precision of early sensorimotor processing.

## Introduction

Our movements are ultimately motivated by the desire to derive rewards and avoid punishments. This is obvious when we move to obtain an immediate reward, such as when reaching for a tasty treat, but also when we move to obtain information that allows us to predict reward availability for future exploitation, such as when we open the refrigerator to plan a future meal. Unsurprisingly, the characteristics of our movements are strongly affected by the prospect of reward. We move earlier, faster, and more accurately towards targets associated with higher rewards (Carpenter and Williams, 1995; Milstein and Dorris, 2007; Manohar et al., 2015; Reuter et al., 2018b; Summerside et al., 2018; Carroll et al., 2019; Galaro et al., 2019). The tendency for more vigorous movement towards rewarding targets makes sense in a dynamic world; if you are slow to grab the treat, somebody else might eat it first.

Despite robust effects of reward on motor behaviour, much of the research on the neural processes that underlie behavioural effects of reward focused on economic decision making tasks (e.g. “bandit” tasks; (Daw et al., 2006; Niv et al., 2012)) or perceptual tasks with trivial motor components (e.g., (Liston and Stone, 2008; Pleger et al., 2008; Serences and Saproo, 2010; Noorbaloochi et al., 2015)). Here we address the situation in which people must rapidly identify the location of a unique and unambiguous visual stimulus, and produce a limb movement toward that location as quickly as possible. The time pressure inherent in this task challenges the core components of sensorimotor control, namely the transformation of sensory inputs about the body and the environment into appropriate motor commands to muscles, while minimising the challenge to perceptual and decision making aspects of goal-directed behaviour. The aim is to establish how sensorimotor processing triggered by target presentation is influenced by the prospect of reward. This would extend evidence that the expected value of actions biases preparatory motor activity towards more frequently required and more highly rewarded actions prior to target presentation (Reuter et al., 2018b).

When actions are stipulated externally, for example when a tasty treat consistently appears in one location and a boring staple consistently appears in another location, the long-run expected value of a potential movement to each location can be calculated as the probability that the movement will be cued, multiplied by the magnitude of its likely reward outcome. Thus, prior to target presentation, both the history of previous actions and the history of previous rewards define expected value. Although both of these factors bias motor preparation when a quick response will soon be required to a yet-unspecified target (Reuter et al., 2018b), there is reason to suspect that they may have different effects on sensorimotor processing once the target has been presented. Behaviourally, we recently reported that although movements were initiated earlier towards both more highly rewarded and more frequently presented targets, movement vigour (i.e., the rate of force production) was enhanced only for movements with a history of yielding high reward magnitude (Reuter et al., 2018b). Moreover, in perception and decision making tasks, conventional indices of neural stimulus processing (i.e., EEG event-related potentials - ERPs) tend to be exaggerated for rewarded versus neutral stimuli (e.g., (Hickey et al., 2010; Meadows et al., 2016; Glazer et al., 2018), whereas ERPs are generally weaker in amplitude for stimuli that are repeated more frequently (e.g., (Maffei et al., 1973; Squires et al., 1975; Movshon and Lennie, 1979; Albrecht et al., 1984)).

Although simple ERP measures of neural dynamics suggest differences between reward and probability responses in some contexts, a more important question is whether the brain develops an enhanced representation of more valuable target locations and associated movement plans. Indeed, multivariate analyses of fMRI data show that the perceptual features of highly rewarded targets are represented more precisely (e.g. (Serences and Saproo, 2010)). Importantly, enhanced neural representation of stimulus features was also reported for expected versus unexpected stimuli, despite the typical attenuation of ERPs for expected stimuli (Kok et al., 2012; Tang et al., 2018). These reports indicate a consistent trend, in which the neural representation of the features of more valuable stimuli is enhanced relative to other stimulus options. Here we address the question of whether both frequency and magnitude components of expected value estimations sharpen post-target sensorimotor processing.

## Methods

### Participants

Forty healthy participants were recruited via the University of Queensland School of Psychology Research Participation Scheme and received either course credit or 60AUD for their time, in addition to and rewards earned during experiments (see below). All participants self-reported to be right-handed, to be free of neurological disorders and recent upper body injuries, and to have normal or corrected-to-normal vision. The study was approved by the ethics committee of the University of Queensland and procedures performed in this study were in accordance with the 1964 Declaration of Helsinki. Participants gave written informed consent prior to taking part in the experiment.

Participants were randomly allocated to one of two experimental groups (N = 20 each). In Experiment 1, one of four targets was associated with a greater reward. In Experiment 2, one target appeared more frequently than the other three. Due to poor EEG signal quality, we excluded three participants from analysis in Experiment 1, and one participant from Experiment 2. Final samples consisted of 17 subjects (10 female, age 21.6± SD 4.1) in Experiment 1 and 19 subjects (10 female, age 24.8 ± SD 7.2) in Experiment 2. Within each experimental sample participants were randomly assigned to either a left or right context group, where the relative reward or probability of a left or right target (respectively) was manipulated.

### Procedure

All participants completed a reaching task and a visual working memory task while electroencephalograms (EEG) and electrooculograms (EOG) were recorded. The reaching task consisted of a baseline phase and a context phase for both experiments. In the baseline phase all targets were equally likely and equally rewarded. In the context phase we manipulated either the reward associated with a certain target (Experiment 1) or the probability of target presentation (Experiment 2). Participants were explicitly told about the probability and reward manipulations prior to the context phase. Each phase consisted of four blocks per task (i.e., four reaching task blocks and four visual detection task blocks in alternating order). Baseline reaching blocks consisted of 120 trials. Context blocks consisted of 120 trials for Experiment 1 and 144 trials for Experiment 2. Each block of the visual task consisted of 192 trials, and was kept constant throughout the session (i.e., reward and probability manipulations were applied only to the reaching task). Prior to the first block, participants completed practice runs of 24 trials each for the movement and vision tasks, which were repeated if participants required additional practice. Testing was completed within 2.5 hours, including time to read and sign the information and consent forms, EEG and EOG setup, and the experimental testing. Participants were offered a break after each block and informed that they could take additional breaks at any time.

### Experimental Tasks and Data Analysis

*Reaching task.* Participants made reaches in the horizontal plane with their right arm while grasping the handle of a two-dimensional planar robotic manipulandum; the vBOT (for full detail of the apparatus see (Howard et al., 2009b)). Visual feedback was provided using a 27” LCD computer monitor (ASUS, VG278H) running at 120Hz mounted above the vBOT and projected to the participant via a mirror. The display was calibrated so that visual feedback appeared in the plane of the limb, and the appearance of a cursor veridically coincided with the physical hand position. The mirror prevented vision of the manipulandum and the participant’s arm. The participant’s arm rested on an air-sled that allowed for near frictionless movement over the table while relaxing their shoulder muscles.

Each trial started with participants moving the robot handle to the start position (circle, radius = 0.5 cm) at the centre of the screen and fixating there. After a variable delay time (500-600 ms), a large target appeared in the periphery (radius = 4 cm), and participants had to make a quick and accurate center-out reach to land the cursor within the target. Targets were presented at four different locations at 90 degree offsets (see Figure 1) in pseudorandomised order, i.e., each target appeared 6 times within 24 trials. The robot enforced a straight-line path between the start and the target position with a mechanical force channel (Scheidt et al., 2000). The mechanical channel ensured that participants accurately reached the target, as long as they moved the target distance of 12 cm ± 4 cm, and prevented differences in success rates between targets. Once participants stopped their movement inside the target circle a short beep was played and they received a feedback message (800 ms duration) about the points they had earned in this trial, and about the cumulative points they had gained within the block. Once the feedback message turned off the robot assisted them to return to the centre.

**Figure 1.**
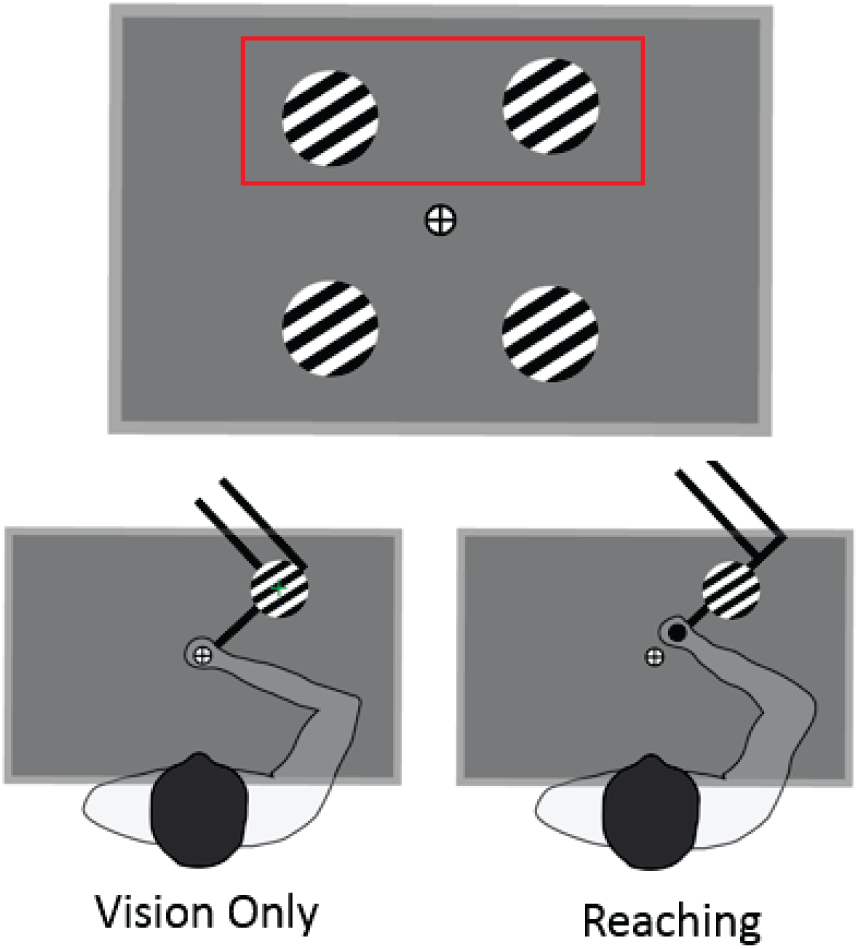
Schematic display of target locations and set up. **A)** Visual targets (high contrast alternating lines in a circular aperture) were shown at four target locations. Either the upper-left or upper-right target (highlighted by the red rectangle) was selected to be associated with a higher reward magnitude or presentation probability. All statistical analyses compared measures between these two upper visual field targets. **B)** In the vision-only task, participants kept their hand and fixation at the centre and counted the number of times a green cross appeared on the screen. **C)** During the reaching task, participants fixated at the centre and reached towards the targets.

Target stimuli were circles filled with a black and white alternating grating. Each of six different grating orientations (in 30° increments) was presented once for each of the four target locations every 24 trials. The targets included this orientation factor to provide an opportunity to assess how value influences the neural representation of visual target features that are irrelevant to the task. Although initial pilot testing suggested that the orientation of the gratings within target stimuli could be decoded from EEG recordings, grating orientation could not be reliably decoded from our experimental data. Consequently, we did not include the orientation of the target stimulus as a factor in any analyses.

In the baseline phase of the experiment, all targets were presented with equal probability and were equally rewarded when successfully reached (1 cent per hit). In the context phase, either the upper left or the upper right target (counterbalanced between participants) was associated with a higher reward magnitude (10 cents per hit, Experiment 1) or it appeared more frequently (i.e., probability of the more frequent target was 50%, instead of ∼17% for the remaining three targets). In addition to the greater monetary reward in Experiment 1, successfully reaching the more highly rewarded target was also followed by a high-pitch ‘bing’ sound (chosen to sound pleasant) instead of a standard mid-pitch beep.

Participants were required to fixate the home circle in the centre of the screen during each trial. If fixation was lost (see below), the trial was stopped and participants received a warning message, reminding them to fixate at the centre. These trials were excluded from all further analyses. To motivate participants to respond as quickly as possible, we set a reaction time limit of 300ms during the practice block. If reaction time exceeded 300ms we terminated the trial and displayed a “Too slow” feedback message. We instructed participants that this reaction time criterion would apply for the entire experiment, whereas in reality, the reaction time cutoff was 600 ms for the baseline and context phases of the experiment.

However, we also presented a sham feedback message that reaction time was “Too slow” on a random of 5% of trials, irrespective of the actual reaction time. These trials were also excluded from all further analyses. We used a relatively lax “true” reaction time criterion that should have been achievable unless there was a lapse of concentration and provided sham “Too slow” feedback to provide an equal incentive to move quickly and an equal success rate for all participants.

Behavioural data were analysed with MATLAB R2017b (MathWorks Inc.). A 5^th^-order Butterworth filter with a low-pass cut-off frequency of 50 Hz was applied to position and velocity data prior to analysis. We analysed reaction time, peak velocity, movement time, and movement amplitude. The movement initiation was measured by applying Teasdale et al.’s (1993) algorithm to the velocity profile for each movement, with a threshold of 10% of the maximum tangential speed. Amplitude was defined as the maximal distance reached from the start location, and movement time was defined as the time between movement initiation and when the maximal distance was reached.

Trials in which reaction times were less than 100 ms were excluded from all further behavioural and EEG data analyses. We also excluded the first 24 trials in the first context block from all further analyses, to ensure that participants had gained experience with the new context. On average, analyses of the baseline and context trials in Experiment 1 included 407 (SD = 39) and 390 (SD = 28) of 480 trials. Similar average retention rates occurred in Experiment 2, with 390 (SD = 51) of 480 baseline trials included in analyses, and 395 (SD = 82) of 576 context trials included.

#### Vision task

In the visual detection task, participants were required to count the number of times they saw a green cross that was occasionally presented either in one of the targets or at the fixation cross. They had to grasp the robot handle, hold their hand in the centre position, and maintain fixation at the centre while a series of visual targets appeared. Target locations were identical to those in the reaching task. Each trial started with a random delay of between 500 and 600 ms before one of the targets appeared for 350 ms. Green cross targets were presented in approximately 20% of trials, such that one cross appeared in each of the four target locations within 24 consecutive trials and at the centre in every 24^th^ trial, unless this trial was randomly selected to have cross at a target location. Participants were asked to count these crosses, to ensure they had attended to the target presentations.

### EOG recording

We recorded EOG via two self-adhesive Ag/Ag electrodes positioned lateral to the eyes. A grounding electrode was placed on the forehead, slightly above the nasion. The signal was amplified with a Grass P511 amplifier (Grass Instruments/AstroMed, West Warwick, RI, USA) and band-pass filtered (0.1 Hz to 300 Hz). At the beginning of the session, prior to the practice trials, participants performed 10 centre-out eye movements to the four target locations. We smoothed the EOG data using a 50 ms moving average and determined peak, rectified EOG amplitudes in each trial. We then averaged the peak amplitudes between trials. Fifty percent of the peak amplitude was set as the EOG threshold criterion, i.e., if the smoothed, rectified EOG amplitude in either the visual or the reaching tasks exceeded the threshold value, the trial was stopped and a feedback message was displayed reminding the participant to fixate.

### EEG recording and analysis

EEG data were recorded using a 64-channel active electrode system (actiCHamp, Brain Products, Munich, Germany). Electrodes were positioned according to the extended 10-20 system (Jasper, 1958). The signal was acquired with a sampling rate of 2500 Hz and was low-pass filtered at 100 Hz. Offline analyses of the EEG data were performed using Brain Vision Analyzer Software 2.0 (Brain Products, Munich, Germany) and MATLAB (MathWorks Inc.).

#### Event-related potential analysis

We pre-processed raw EEG data using a method similar to that described in (Reuter et al., 2018a). In brief, we used an average reference, down-sampled to 1000 Hz, and applied a high-pass filter of 0.1 Hz, a low-pass filter of 30 Hz, and a notch filter around 50 Hz. For the reaching task, data were segmented into 1 second epochs (−200 to 800 ms relative to target appearance). Data collected during the vision-only task were partitioned into 700 ms epochs (from −100 to 600 ms relative to stimulus onset). Epochs were baseline corrected using the pre-stimulus period. Blinks were identified using the ocular artefact removal algorithm included in Brain Vision Analyser, and trials with blinks were excluded from all further analyses. EEG activity with voltage magnitudes exceeding 100 µV, or voltage changes of more than 50 µV in a 100 ms time-window, were rejected channel-wise as artefacts. Finally, we computed event-related potentials for each target location by averaging the relevant pre-processed epochs.

We analysed P1 and N1 components, which are thought to be related to early visual information processing ((Hillyard and Anllo-Vento, 1998; Di Russo et al., 2003). N1 and P1 ERP components were estimated at electrodes P07 and P08, situated over the occipital-parietal lobe in each hemisphere (Jasper, 1958). We first identified the timing of the largest positive/negative peak within pre-defined timing windows for each component (100 to 200 ms after stimulus onset for the P1, and 150 to 250 ms following stimulus onset for the N1), and then computed the average of ERP data within a 50 ms window centred at that peak-timing. Note that we analysed component amplitudes at occipital-parietal sensors contralateral to the hemifield of the visual stimuli.

Statistical analyses for reaction times, peak velocities, movement amplitudes and P1 and N1 amplitudes were conducted separately for Experiments 1 and 2 using SPSS for Windows version 25.0 (IBM Corp., Armonk, NY). We used a 2 (Value: baseline phase, context phase) × 2 (Target: less valuable, more valuable) within-subjects ANOVA model and applied the Huynh–Feldt non-sphericity correction to within-subject comparisons where appropriate. Effect sizes are given as partial eta-squares (ηp^2^).

To provide a general overview of the EEG responses to the movement and vision only tasks, we also show ERP results for five midline electrodes (Fz, FCz, Cz, CPz, Pz). Average ERPs time-locked to the target presentation and the movement initiation time were constructed for left and right upper visual field target locations for each participant. To assess whether reward or frequency manipulations influenced ERP amplitudes, we compared the average ERP amplitude in sequential 75ms bins (with 25ms overlaps between consecutive bins) between the manipulated and control targets with Bonferroni-corrected t-tests via MATLAB (MathWorks Inc).

### EEG decoding method and analysis

We used a multivariate backward linear decoding method to estimate the precision and accuracy of target location information contained in the EEG recording at a range of time intervals around the time of stimulus presentation. Our approach is similar to forward encoding model approaches used previously (Brouwer and Heeger, 2009; Tang et al., 2018; Chen et al., 2021; Foster et al., 2021). However, we used a backward (decoding) rather than forward (encoding) model as we found this approach more robust to noise in the EEG recording during pilot testing. This method does not attempt to directly decode the categorical label of the target location, or the discrete angle of target presentation. Instead, we assumed that the EEG signal at each electrode reflects the linear sum of eight spatially selective channels tuned for target or movement directions distributed equally around 360 degrees. By identifying a weight matrix that defines a linear mapping from EEG electrode space to channel space from a set of training trials, we could estimate the weighted contribution of each channel direction to the EEG signal amplitude from each trial. From these channel response amplitudes, we could derive measures of the selectivity of the neural representation of target or movement directions at each time point during the course of each trial.

#### Decoding method

First, we represented target locations in polar coordinates. We discarded radius coordinates as all targets were the same distance from fixation, yielding a one-dimensional coordinate for each target location. We then defined a set of eight spatially-selective channels that are each maximally responsive to a given angle, with tuning curves defined by a raised-power cosine function. We modelled the response amplitude of each channel, *c*, with a pre-defined mean, *µ*, as a function of the target location, *θ*, on each trial, *t* (see Equation 1). These channels were spaced uniformly around the circle such that the sum of activity from this set of channels was approximately equal for every point on the modelled circle. This approach allows for any number of channels to be modelled and subsequently decoded, but the specificity of each channel increases with channel-density. Therefore, a high number of channels produces high redundancy (as many channels would never be active, given we had only four target locations), whereas a low number of channels limits the resolution of the reconstructed spatial tuning curves. To balance these considerations, we selected eight channels for our model.

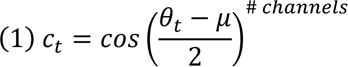

Since some trials were removed from analyses during pre-processing, we often had a slightly unbalanced number of trials for each target location. To ensure unbiased decoding, we selected (at random, without replacement) an equal number of trials at each target location from the set of available trials. This yielded *T* trials, which differed slightly between subjects and experiments, with more trials were discarded in Experiment 2 (as one target location was presented much more frequently, by design).

With the spatially-selective channels defined, we then computed the required amplitude of the response of each modelled channel given the target location on each trial. This yielded a set of channel responses, *C*, with dimension *T* × 8, where *T* is the number of trials being analysed. The goal of the decoding approach was to find an optimal set of weights, *W*, that could extract an accurate estimate of the response amplitude of the modelled spatially-selective channels. To do this, we first solved for *W* in Equation 2 (below), using training-case EEG data, *E_1_*, and modelled channels-responses, *C_1_* (see Equation 3). We then applied the trained weight matrix, *W*, to independent EEG data from each trial of the experiment, *E_2_*, allowing us to extract an estimate of the response amplitude on each spatially-selective channel of *C_2_* (see Equation 4).

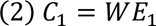

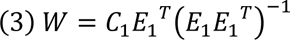

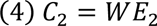

In this approach, the input EEG is a matrix, *E*, containing the average EEG data within a pre-defined time-window at each scalp channel, on each trial (i.e., a matrix of dimension *T* × number of scalp channels). For convenience, we included a third-dimension to this matrix that spanned the time-windows we had pre-defined (i.e., *T* × number of scalp channels × number of time-windows). To solve for *W*, we first isolated one trial for each of the four target locations and used the remaining trials as training data. We then computed the average EEG activity within 16 ms time-windows, with each window offset from the preceding window by 4 ms (a total of 247 windows, spanning −200 ms to 800 ms relative to stimulus onset). We then solved for *W* for each time-window using Equation 3, and applied the solved *W* for each time-window to the isolated test-case EEG data using Equation 4. We repeated this process for every set of four test-case EEG trials, comprising one instance of each target location. This yielded a set of channel response amplitudes, *C_2_*, for every trial at every specified time-window.

The outcome matrix, *C_2_*, contains estimates of the response amplitude on each of the spatially-selective channels. Since we had eight channels and four target locations, we had a specific channel that corresponded to the target location on each trial. We circularly shifted the decoded responses on each trial such that the channel that ought to be maximally responsive was aligned across trials. This yielded a matrix of decoded spatial-tuning curves, independent of target-location between trials.

#### Analysis of decoded tuning curves

We first applied the decoding method described above to EEG data recorded from all analysed participants (participants excluded from ERP analyses were also excluded here), in all active movement conditions. We then compared, within-subjects, the decoded spatial-tuning curves between the high and low value targets in the upper-visual-field. To do this, we computed a measure of the centrality of the channel responses for every trial in every time-window, by subtracting the mean response amplitude on all non-target channels from the response amplitude of the target channel. This controls for the base amplitude of decoded responses for each participant, providing an estimate of the response of the target channel over and above the response of all non-target channels. Statistical analyses comprised comparisons of the relative centrality of decoded responses for the value-manipulated target versus the upper-visual-field control target location, in both the baseline and value-manipulation conditions.

To ensure decoding was successful, we first collated and analysed the relative centrality of decoded responses from all trials in the baseline and value-manipulation conditions. If decoding was accurate and precise, the relative centrality of the decoded responses should reliably differ from zero. We used a series of within-participant *t*-tests to compare participants’ overall mean relative centrality to zero at each time-window, with an alpha-criterion Bonferroni-corrected for 247 comparisons (Benjamini and Hochberg, 1995; Storey, 2002). We found evidence that our decoding method was able to accurately and precisely decode the location of visual targets soon after target presentation. We used a similar analytic approach to estimate the effect of our value-manipulations on target-location decoding for final analyses. However, we used a permutation test (10,000 permutations) for final analyses (Maris and Oostenveld, 2007) rather than within-participant *t*-tests, comparing the manipulated versus control upper-visual-field target location in the baseline condition, and again in the value manipulation condition.

## Results

We conducted two experiments to determine the effect of manipulating the expected value of reaching movements to specific directions. In our first experiment, we manipulated the reward distribution; one of the four targets was more highly rewarded during the reaching task, although all four targets were equally probable. In our second experiment we manipulated target probability; one target appeared three times more frequently than each of the other three targets during the reaching task, although movement to each target incurred an equal reward. We also established the influence of exposure to reward and probability manipulations in the reaching task upon neural processing during an alternative task, in which participants did not move and merely counted occasional events while visual stimuli were presented at the same target locations used in the reaching task. We used a multivariate EEG linear decoding approach to measure the precision of representation of target location (and/or movement direction) in both reaching and non-reaching tasks, although the valuable stimulus attribute (direction) was only behaviourally relevant during reaching trials.

### Behavioural results

We analysed reaction times, peak velocities, and movement amplitudes as a function of target position. We restricted all analyses to the upper two targets to avoid dramatic differences in the cortical topography of early visual processing between the upper and lower visual fields. In the baseline condition, these two targets had the same value (i.e., same associated reward and probability of occurring). In the context condition, one of these targets was associated with a higher expected value, either by manipulating its reward (Experiment 1) or its probability of occurring (Experiment 2).

Grand mean reaction times at baseline were 225 ± 18 ms in Experiment 1, and 218 ± 22 ms in Experiment 2. These reaction times are quite short for a four-choice manual response task, and suggest that our manipulation of response time feedback to encourage rapid responses were successful in promoting early responses. Peak movement velocities were 50 ± 13 cm/s in Experiment 1, and 45 ± 7 cm/s in Experiment 2. Movement amplitudes were 11.7 ± 0.5 cm in Experiment 1, and 11.7 ± 0.7 cm in Experiment 2. Movement durations were 458 ± 90 ms in Experiment 1, and 510 ± 112 ms in Experiment 2. We found no significant differences between the two upper visual field target locations on any of these measures at baseline in either experiment.

We used a 2 (Block: Baseline, Context) × 2 (Target: less valuable target, more valuable target) within-subjects ANOVA to assess the influence of target value on participants’ behaviour. In both experiments, participants initiated their responses earlier to the more valuable targets, evidenced by significant interactions between Block and Target factors (Experiment 1: *F*(1,18) = 27.4, *p* < 0.001, *n*^2^_p_ = 0.60; Experiment 2: *F*(1,18) = 30.6,*p* < 0.001, *n*^2^_p_ = 0.63; see Figure 2). These results indicate that our manipulations of value were effective in modulating participants’ behaviour toward these targets. Participants’ responses to a more rewarding target in Experiment 1 were also performed with a higher peak velocity, (Block × Target: F(1,18) = 16.9,p = 0.001, n^2^_p_ = 0.53) and had a larger amplitude (Block × Target F(1,18) = 9.6,p = 0.007, n^2^_p_ = 0.38). Block by target interactions for peak velocity (p = 0.63) and movement amplitude (p = 0.27) were not significant for Experiment 2, where one target appeared more frequently. Thus, while both reward and frequency shortened reaction times, movement velocity and distance were modulated only by reward and not by frequency.

**Figure 2.**
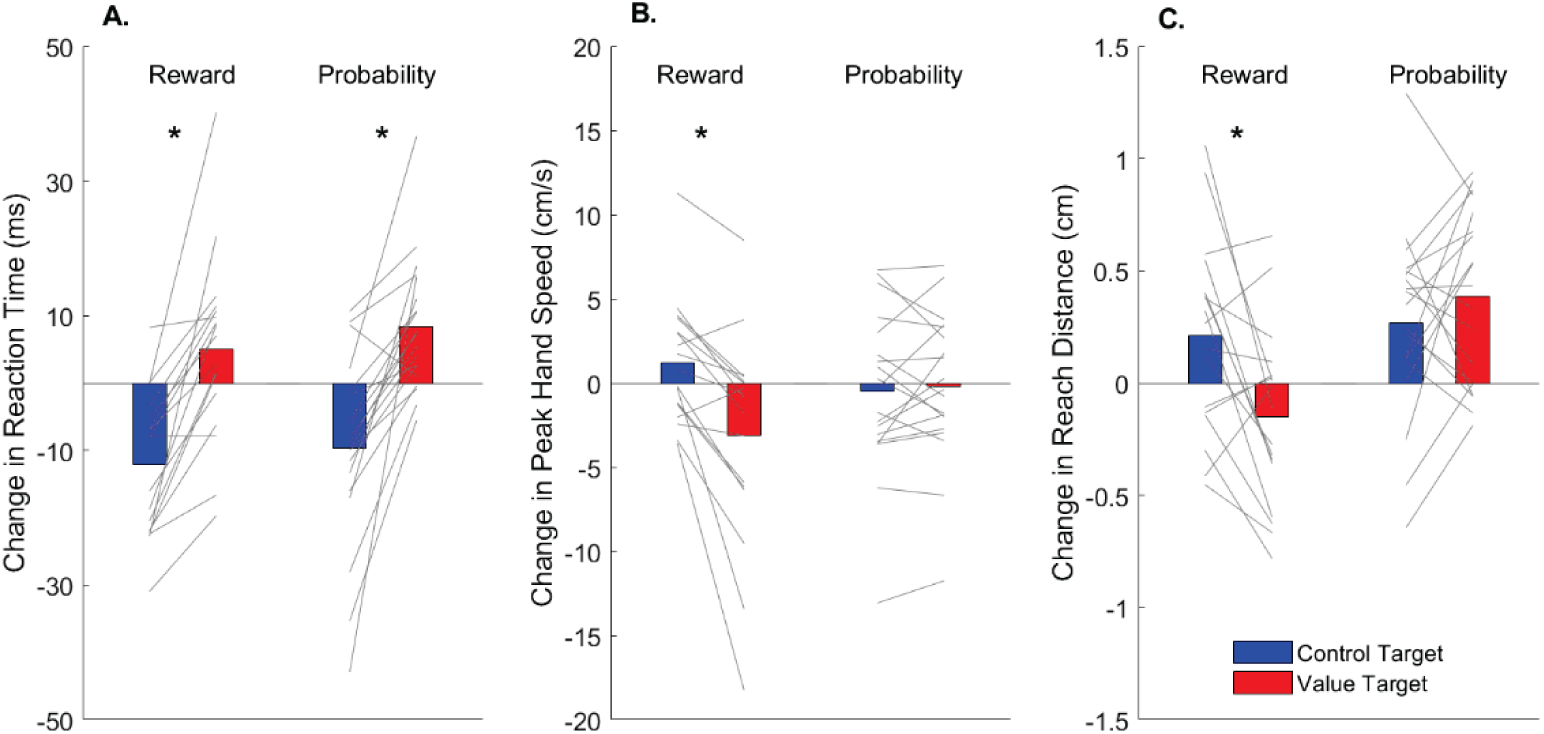
Changes in reaction times, peak velocity, and movement distance as a function of target position (blue = control target position, red = valuable target position) for the Reward (experiment 1) and Probability (experiment 2) experiments. Each line shows the changes from baseline (baseline minus context) for the movements of an individual participant to each of the targets, and the coloured bars show the mean values for the groups. Asterisks indicate significant ANOVA target by block interaction effects.

### Classic P1/N1 ERP results

We first checked if classical ERP components associated with early visuospatial processing (Hillyard and Anllo-Vento, 1998; Di Russo et al., 2003) were affected by our value manipulations. Figure 3 shows grand average P1 and N1 ERPs for both conditions and experiments. Initial inspection of the spatiotemporal distribution of average ERPs indicated that P1 and N1 peak latencies were earlier, and N1 amplitudes larger, at electrodes contralateral to the visual stimuli.

**Figure 3.**
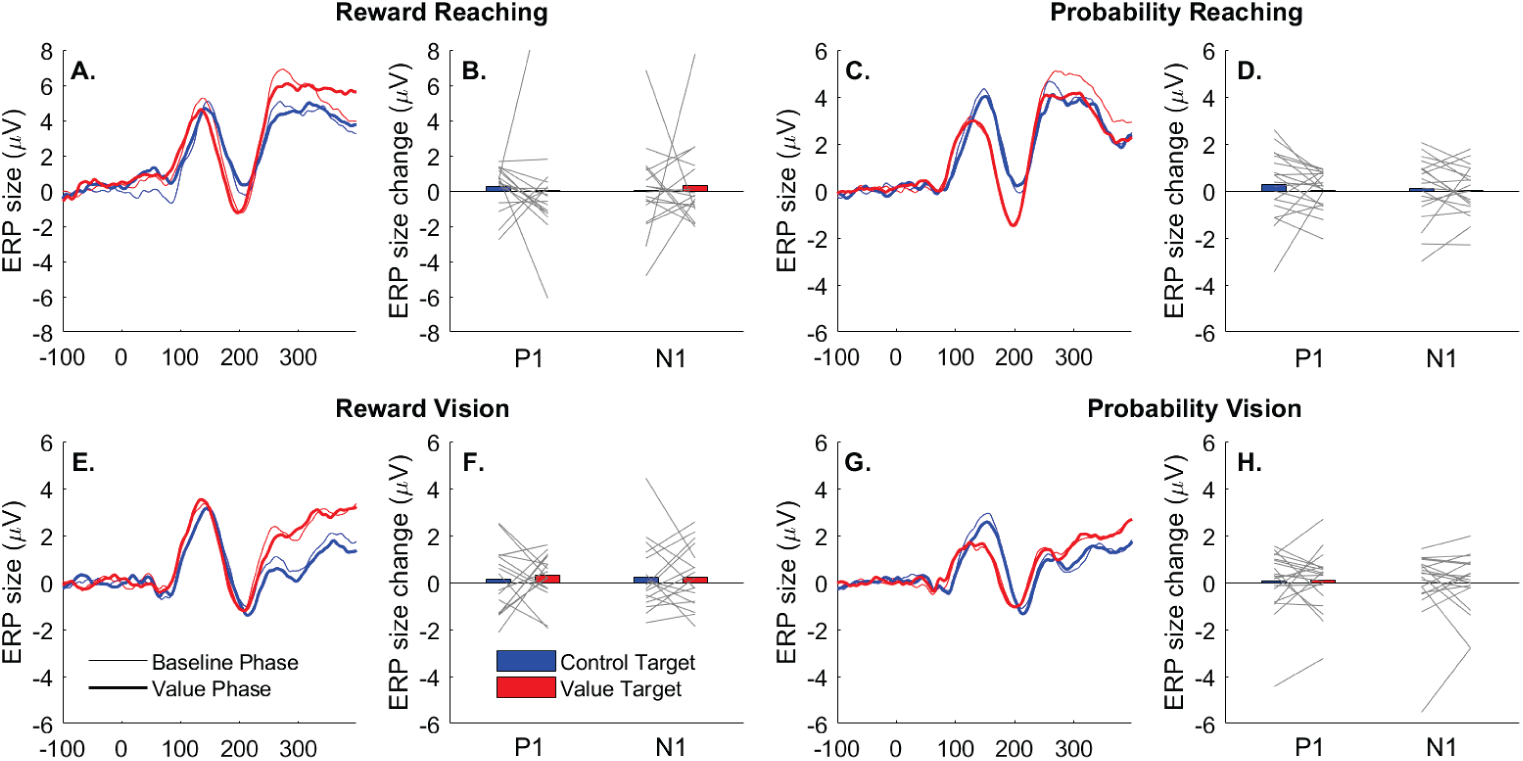
Grand average ERPs at electrodes contralateral to the target (Po7 or Po8) are shown for targets presented during the reaching task in panels A and C, and during the visual detection task in panels E and G. Thin traces show ERPs extracted from the baseline condition, and thick lines are ERPs from the blocks in which we manipulated target value. Red lines denote ERPs evoked by the valuable target, and blue lines denote traces to the control target. The left panels show results for Experiment 1, where we manipulated the reward associated with one of the target positions, and the right two panels show results for Experiment 2 where we manipulate the probability of target appearance. In panels B, D, F, and H, we show the average change in ERP amplitudes from the baseline to the context condition for control (blue) and valuable (red) targets and for each participant.

To test whether EPRs related to visual processing were affected by the value manipulation during the reaching task, we analysed P1 and N1 mean amplitudes at electrodes contralateral to the target stimuli. For both experiments, and for both P1 and N1 amplitudes, there were no statistically significant main or interaction effects in the Value by Target ANOVAs. There were also no significant main or interaction effects for P1 and N1 amplitudes for the vision only task (all p > 0.47). Thus, there is no evidence from early visuospatial processing ERPs that stimulus processing during the vision-only task was affected by value associations experienced in the reaching task.

### Midline ERP results

To provide a general overview of the EEG responses to the movement and vision only tasks, we show the ERP results for five midline electrodes from frontal to parietal brain regions. Figures 4 and 5 (panel A) show that for both experiments, there is a general pattern of positivity at Pz from approximately 150ms prior to movement onset followed by a second period of positivity from soon after the average time of movement initiation (dotted line at ∼200ms) for ∼300ms. This pattern is inverted for the frontocentral electrodes (Fz, FCz, Cz). In the vision only task, the initial waveforms are similar to those in the movement task for frontocentral electrodes, but delayed or inverted for Pz. The late potentials after 200ms are dramatically reduced.

**Figure 4.**
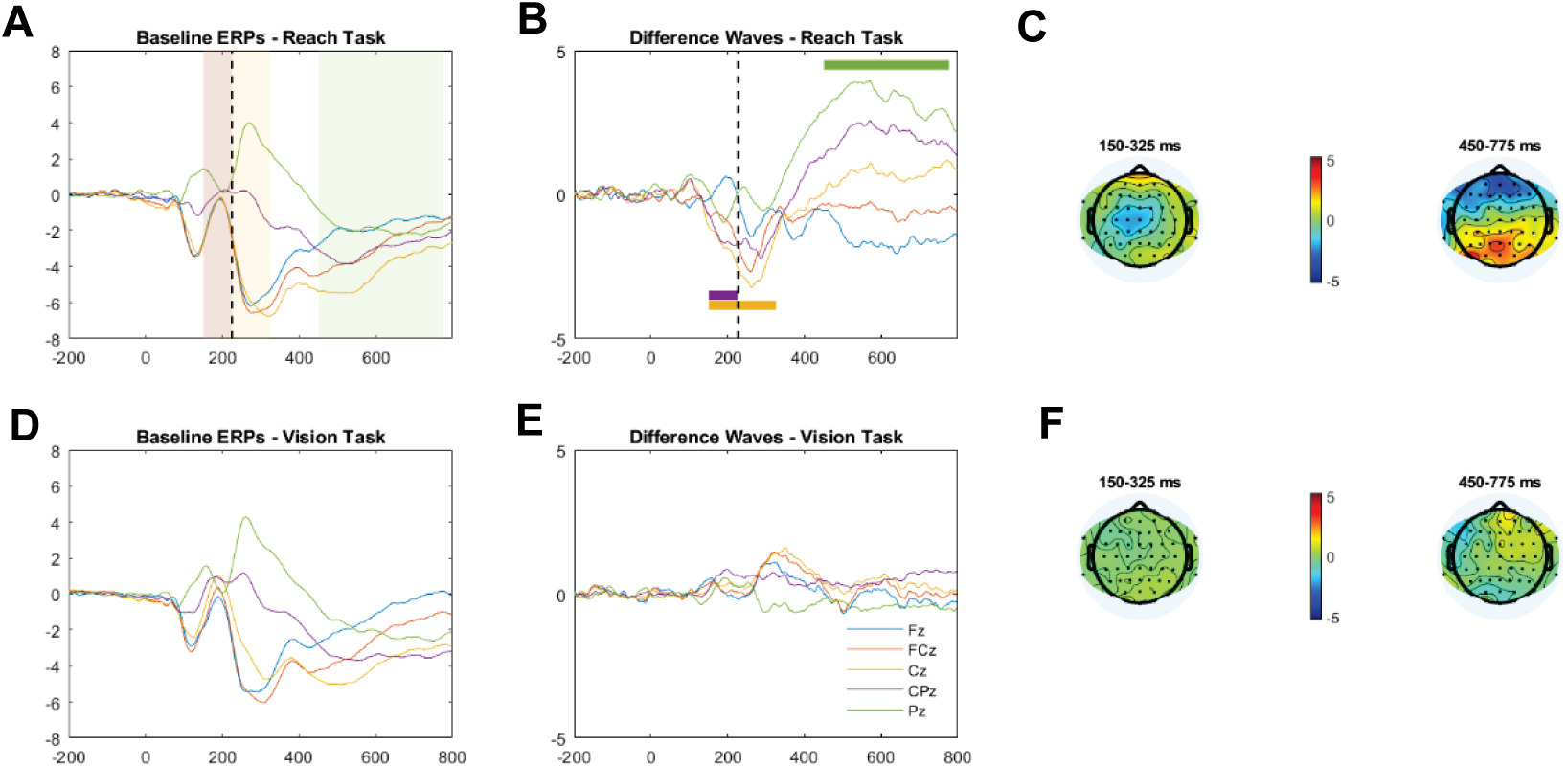
Grand average ERPs time-locked to target presentation and associated difference waves for ERPs between rewarded and control targets in both reaching and vision only tasks. A) & D) show the grand average ERPs at midline electrodes from frontal to parietal regions for the trials to both upper visual field targets in the baseline phase of the reaching (A) and vision only (D) experiments. Time 0 equals target presentation time. Shaded regions in A) and D) illustrate times at which there were significant differences in ERP amplitude between rewarded and control movements as defined in plot B). B) & E) show the difference waves between rewarded movements and control movements. Positive potentials indicate that the ERP for the rewarded movement was more positive than the ERP for the control movement. Coloured bars signify significant differences at specific electrodes according to the colours defined in the legend. C) & F) show heat maps for the difference waves between high value and control movements at early and late times with significant effects.

**Figure 5.**
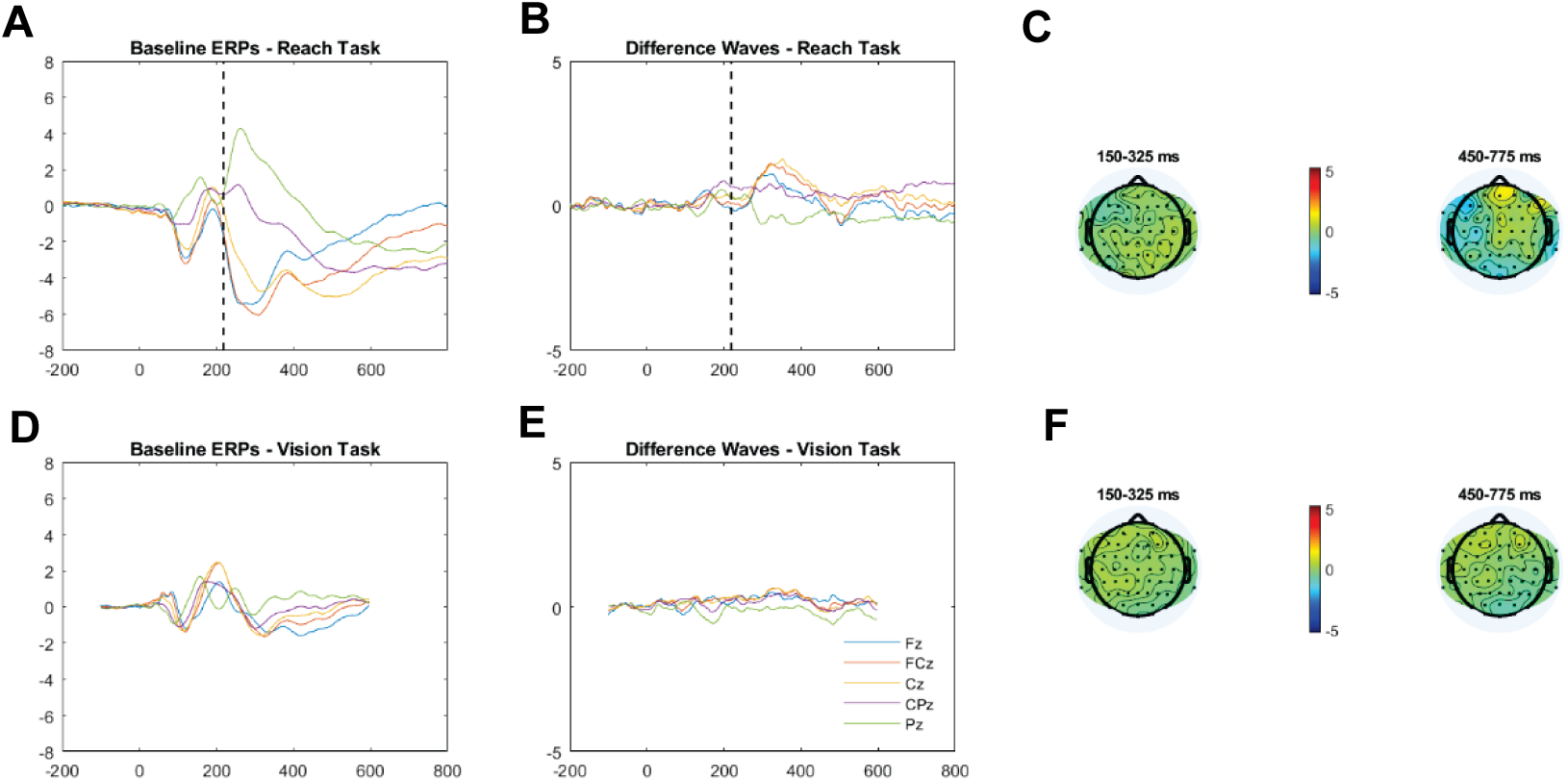
Grand average ERPs time-locked to target presentation and associated difference waves for ERPs between frequent and control targets in both reaching and vision only tasks. A) & D) show the grand average ERPs at midline electrodes from frontal to parietal regions for the trials to both upper visual field targets in the baseline phase of the reaching (A) and vision only (D) experiments. Time 0 equals target presentation time. B) & E) show the difference waves between frequently made movements and control movements (frequent movement ERP – control movement ERP, as above). There were no significant differences at any electrodes. C) & F) show heat maps for the difference waves between high frequency and control movements at the early and late times that had significant effects in the Reward task.

We next checked if the prospect of reward or increased frequency of presentation influenced the amplitude of these midline responses. Panel B of Figure 4 shows the difference wave between EEG potentials evoked by the rewarded target minus the control target. Potentials were significantly more negative for Cz and CPz around 200ms after presentation of the rewarded target than the control target, and more positive for Pz from 450ms to 775ms after target presentation of the rewarded target. The scalp distributions of the difference waves are illustrated in Figure 4C. In terms of the task performance, the timing of these EEG effects coincides with just prior to movement onset into early movement initiation for the Cz and CPz electrodes, and mid-late movement for the Pz electrode. Given the average movement time of ∼450-500ms (plus reaction time of ∼200ms), the Pz effect would appear too early to be related to reward feedback. Importantly, there were no significant differences in ERP amplitudes for rewarded versus control targets in the vision only task, suggesting that the effects relate specifically to sensorimotor processing and/or a reward contingent upon action.

Panel B of Figure 5 shows the difference wave between EEG potentials evoked by the more frequent target minus the control target. Here, there were no significant differences in ERP amplitudes in either the movement or the vision only task, suggesting that repletion of movement or an increased probability of target appearance did not affect these gross ERP measures.

Finally, we considered ERPs time-locked to the movement onset rather than the target appearance for the two experiments (Figure 6). The basic pattern of results is very similar to that of from the target-locked analysis, but the timings with respect to task performance are even clearer. Again we see significant effects from 100ms prior to movement onset into early movement initiation for the Cz and CPz electrodes, and from 250-375ms into the movement for the Pz electrode (i.e. well before reward feedback). Overall, the data show gross ERP amplitude differences between rewarded movements but not more frequently executed movement at midline parietal and central electrodes. The critical next question is whether these gross amplitude changes are accompanied by differences in the precision with which movement directions are represented by the brain.

**Figure 6.**
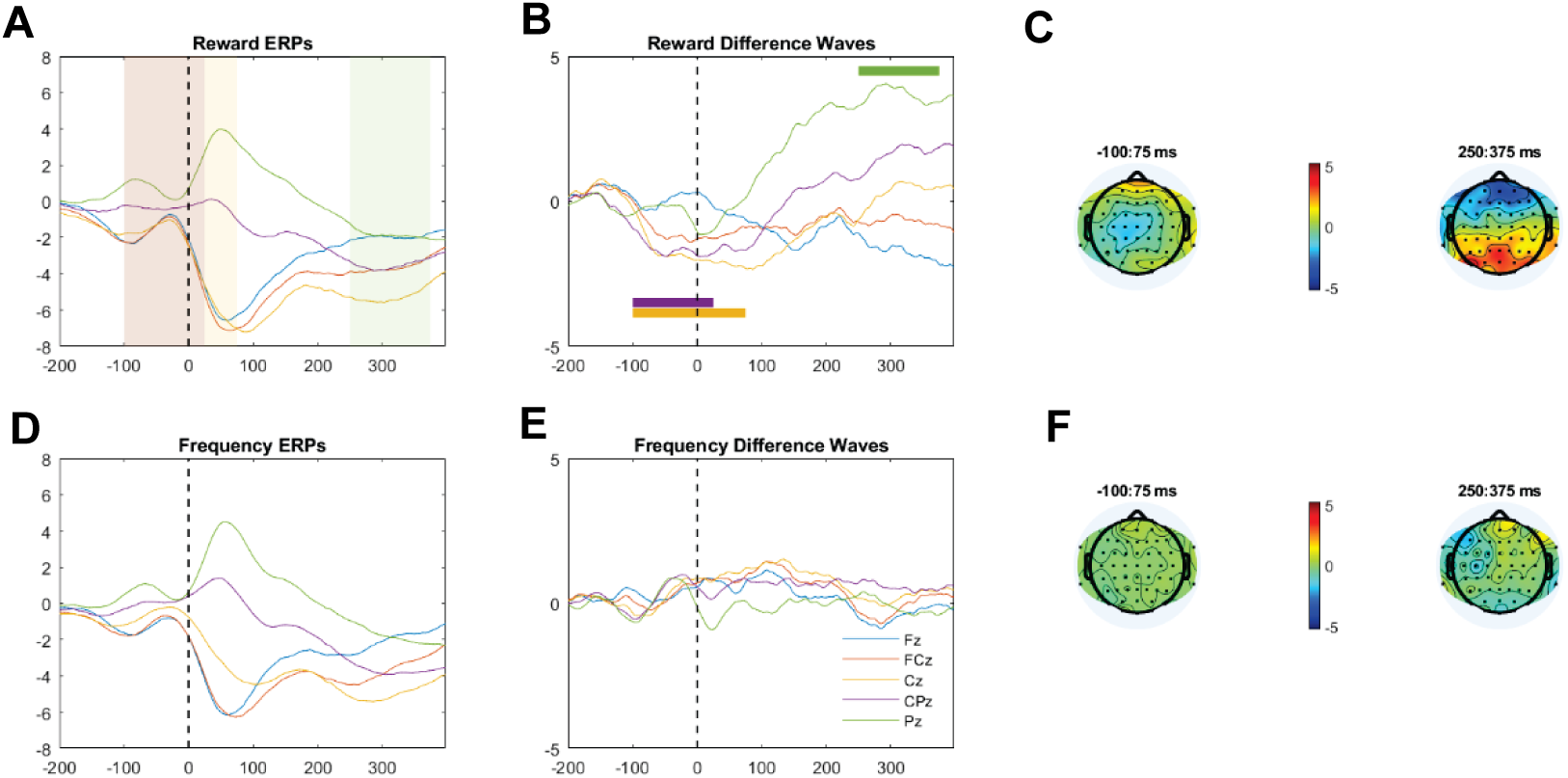
Grand average ERPs time-locked to movement initiation and associated difference waves for ERPs between Rewarded/Frequent and control targets. A) & D) show the grand average ERPs at midline electrodes from frontal to parietal regions for the trials to both upper visual field targets in the baseline phase of the Reward (A) and Frequency (D) experiments. Time 0 equals movement initiation time. Shaded regions in A illustrate times at which there were significant differences in ERP amplitude between rewarded and control movements as defined in plot B). B) & E) show the difference waves between high value movements and control movements (frequent or rewarded movement ERP – control movement ERP, as above). Coloured bars signify significant differences at specific electrodes according to the colours defined in the legend. C) & F) show heat maps for the difference waves between high value and control movements at early and late times with significant effects.

#### Experiment 1 - Decoding results

We first tested whether our analytic method could successfully decode target locations, regardless of the associated value. We used a measure of centrality for decoded responses, allowing us to analyse the extent to which the target channel was responsive over and above non-target channels. We used a series of within-participant *t*-tests to test the null hypothesis that unsuccessful decoding would yield relative centrality values equal to zero. We found a large swathe of time-windows with evidence to reject this null hypothesis, even using a conservative alpha-criterion (0.05, Bonferroni-corrected for 247 comparisons), indicating reliable decoding of the target location. Reliable decoding was observed from the time-window spanning 72–88 ms post-stimulus until the end of the ERP epoch (800 ms after stimulus onset; see Figure 7). Since these results were obtained on a measure of relative centrality, we were confident the decoding method was generally able to provide accurate and precise decoding of the target location.

**Figure 7.**
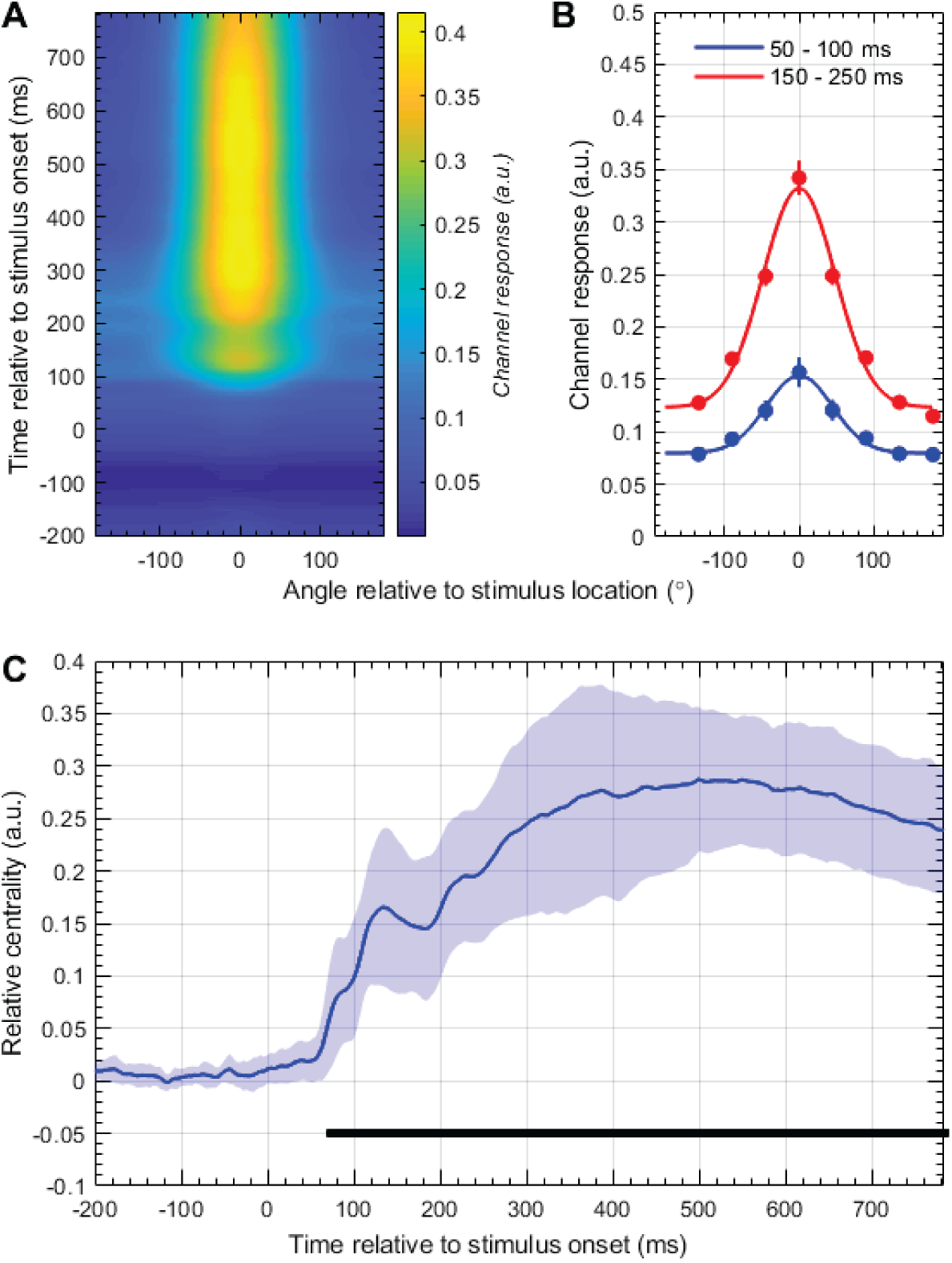
Overall decoding results in Experiment One. We found that the decoding method was able to provide accurate and precise estimates of the target location. **(A)** Gaussian-smoothed heat-map of the response amplitude of decoded spatially-selective channels. Note that decoded outcomes are shifted to align with the physical target location, making tuning curves comparable across trials and conditions. Accurate and precise decoding emerges <100 ms post-stimulus, and is sustained for the duration of the recorded EEG epoch. **(B)** Here we show tuning curves averaged within two periods post-stimulus. Raw values are indicated by markers, with error bars reflecting between-subject standard-error. We have fitted Gaussian functions to these data (solid lines), for illustrative purposes only. Tuning curves are well-captured by Gaussian functions, are accurately positioned over zero degrees (i.e., zero error between decoded and true target location), and are precise. Note the baseline offset of the later-epoch tuning curve, clearly demonstrating the importance of the relative centrality measure used for final analyses. **(C)** Average relative centrality over time, including shaded regions reflecting the between-subject standard-deviation. Positive values indicate accurate and precise decoded spatial-tuning curves. The solid black line indicates time-windows with significantly non-zero relative centrality.

Having found evidence to suggest our decoding method could produce accurate and precise spatial tuning curves, we then compared the decoded tuning curves for individual target locations as a function of their associated value. In the baseline and reward conditions, we compared the relative centrality of decoded spatial tuning curves for the target location associated with higher value (for the baseline block, the target that would later be associated with higher value) with those from the control target location. While both locations were accurately and precisely decoded in the baseline condition, as indicated by the relative centrality values (see Figure 8, Panel A), we found no pattern of reliable differences that would suggest bias in the neural representation of equally valuable target locations. However, a clear and compelling trend emerged when we conducted the same comparison on EEG data recorded during the reward condition (see Figure 8, Panel C). We found sustained and reliable differences in the relative centrality of decoded spatial tuning curves for the rewarded versus control target location in time-windows from 136 ms post-stimulus through to 368 ms post-stimulus. Note however that two time-windows late in this period, from 336 to 340 ms post-stimulus, did not show significant differences. To illustrate the difference in effects between baseline and reward conditions, we have included the tuning curves for the rewarded and control locations averaged over all time-windows that were significant in the reward condition (Figure 8, Panels B and D).

**Figure 8.**
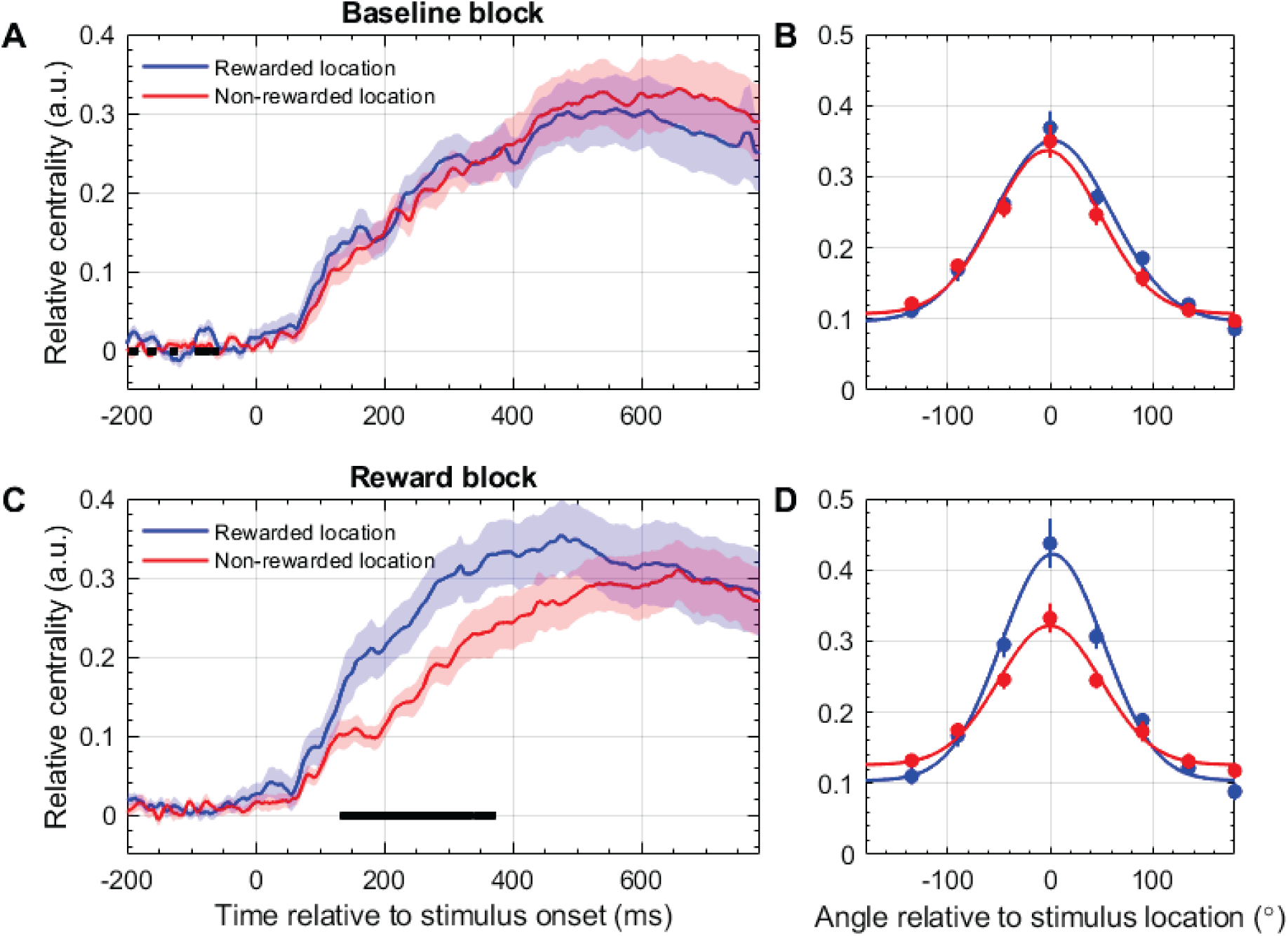
Decoded spatial tuning curves as a function of experimental condition and associated reward in Experiment One. **(A)** Relative centrality of the spatial tuning curves decoded following presentation of the visual target to be rewarded in subsequent experimental blocks and the control target. Shaded regions reflect one between-subjects standard deviation. Black bars indicate time-windows that pass permutation significance testing. There is no reason for the target locations to differ in decoding results during this phase of the experiment. **(B)** Average decoded channel response amplitude during the baseline block, at the time-windows that pass significance testing in the reward block (i.e., Panel C). We fitted a Gaussian function with free mean, amplitude, standard deviation, and offset parameters, for illustrative purposes only. Error bars reflect one between-subjects standard-error of the mean. **(C)** Relative centrality of the decoded spatial tuning curves for the rewarded and control target locations in the Reward phase of the experiment. Plotting conventions are the same as those used in Panel A. We found reliable differences in the relative centrality of decoded spatial tuning curves for rewarded versus non-rewarded stimuli in the range 136 to 368 ms post-stimulus. **(D)** As in Panel B, here we present the decoded spatial tuning curves for the rewarded and control target locations during the time-windows that passed significance testing in Panel C. As suggested by the relative centrality measure, we decoded a greater amplitude, and more precise, spatial tuning curve from the rewarded versus control target location.

#### Experiment 2 - Decoding results

We followed the same analytic procedure for data recorded in Experiment 2 as we had for Experiment 1. We first quantified and tested the overall reliability of the relative centrality of the decoded spatial tuning curves. We found compelling evidence of accurate and precise target location decoding for Experiment 2 (see Figure 9). The pattern of results is very similar to that observed in Experiment 1, with decodability emerging slightly earlier than 100 ms post-stimulus.

**Figure 9.**
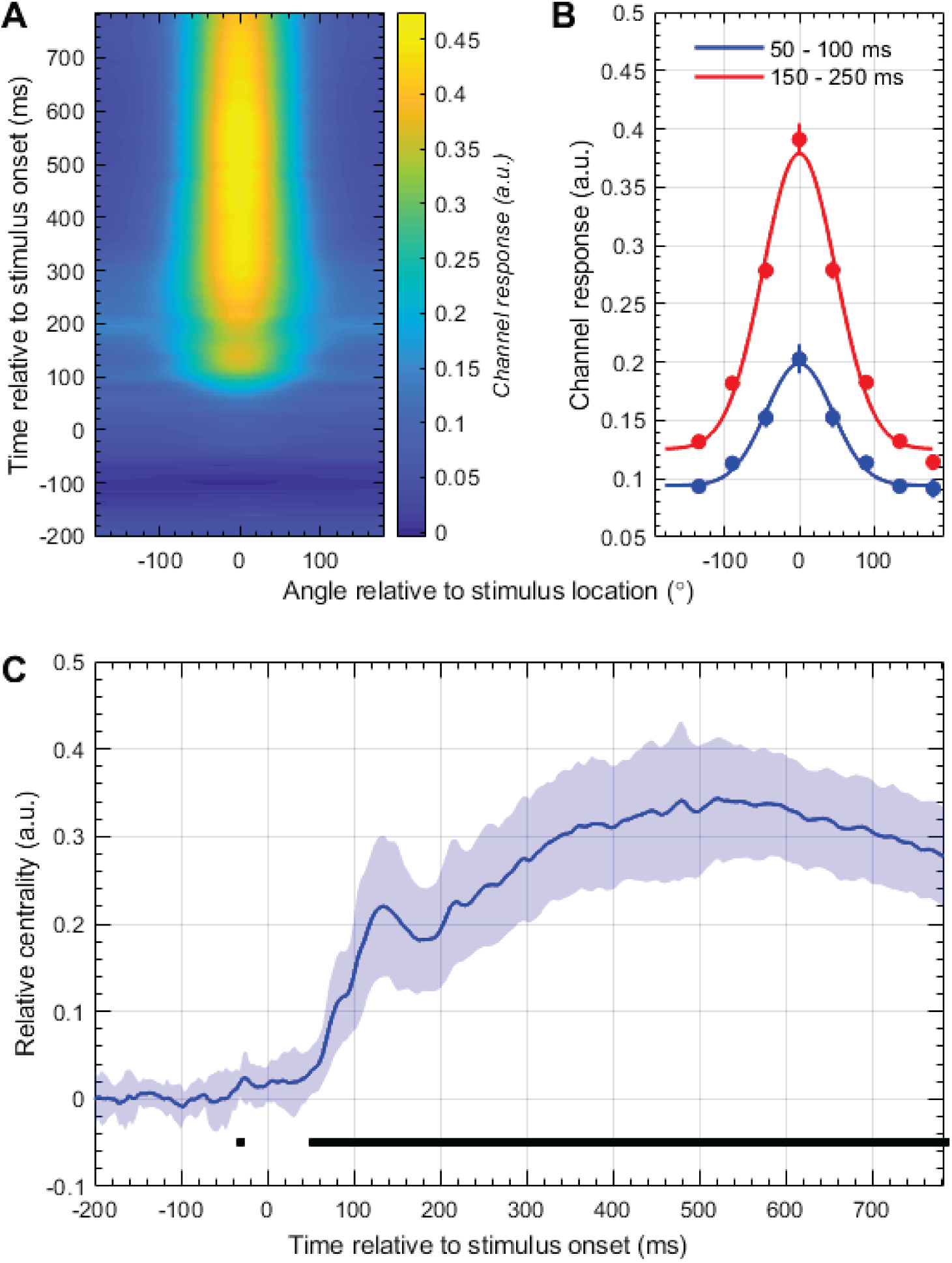
Overall decoding results in Experiment Two. Plotting conventions are the same as those used in Figure 8.

Following the procedure outlined in Experiment 1, we then compared the relative centrality of the decoded spatial tuning curves in the baseline and frequency-manipulation conditions. As in Experiment 1, we found no clear pattern of differences between equally valuable target locations during the baseline condition (see Figure 10, Panels A and B). However, unlike in Experiment 1, we found no clear differences in the relative centrality of decoded spatial tuning curves in the value-manipulation condition, this time a manipulation of the target frequency (See Figure 10, Panels C and D). Together, these results suggest that the neural representation of spatial locations is not modulated by value, which comprises both reward and frequency, but is specifically modulated by the magnitude of the reward associated each spatial location.

**Figure 10.**
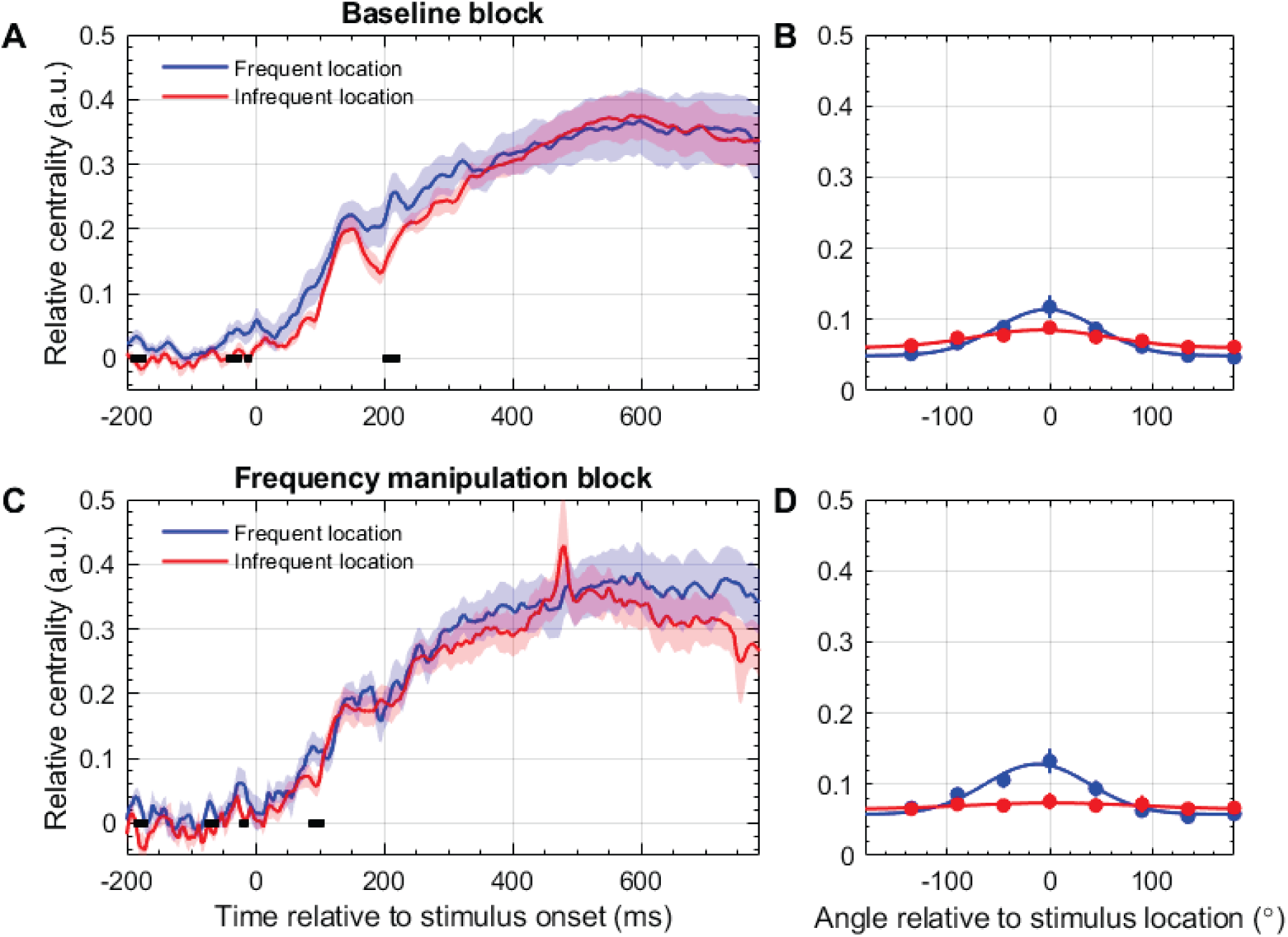
Decoded spatial tuning curves as a function of experimental condition and associated reward in Experiment Two. **(A)** As in Experiment 1, and consistent with our predictions, we found no clear evidence of differences in the relative centrality of spatial tuning curves decoded from target location during the baseline block. **(B)** Spatial tuning curves during the baseline block, at the time-windows that passed significance permutation testing in the frequency-manipulation block. Note that these curves are so shallow because many of the time-windows that passes significance testing in the frequency-manipulation block were pre-stimulus, indicative of false-positive test results. **(C)** Unlike in Experiment 1, and contrary to our predictions, we found no clear pattern of differences in the relative centrality of the decoded spatial tuning curves for the frequent and control target locations. This suggests that neural representations of spatial locations are not modulated by value, but specifically by the reward associated with spatial locations. **(D)** Decoded spatial tuning curves from the frequency-manipulation condition at the time-windows that passed significance testing during the frequency-manipulation condition. As in Panel B, these tuning curves are so shallow because of the timing of the significant windows.

## Discussion

The key finding of this study is that the precision of sensorimotor processing is sharpened when a reaching movement is associated with high reward magnitude, but not when the movement is more frequently repeated. This implies that reward and action history have distinct effects on the way that sensory information that triggers motor behaviour is processed in the brain, even though both reward and action histories should contribute to pre-target estimations of expected value. This distinction in *post-target* processing contrasts with evidence that *pre-target* sensorimotor preparation is enhanced for more probable movements despite delivering equal rewards (Reuter et al., 2018b). There is also strong behavioural evidence that asymmetry in action history leads to pre-target biases in sensorimotor preparation (Marinovic et al., 2017; Mawase et al., 2018; Tsay et al., 2022), since the extent to which movement is biased towards more frequently presented targets is magnified as a function of reduced time available between target presentation and movement initiation. Our current data show that pre-target sensorimotor activity associated with the prediction of upcoming actions does not necessarily lead to a sharpened neural representation of that action after it has been triggered by an incoming sensory signal. By contrast, if the incoming sensory information indicates that the required movement is associated with high reward magnitude, the subsequent neural activity from ∼135ms post-stimulus onset reflects a more precise spatial representation of the target and movement direction.

How might reward sharpen the spatial representation of the action?

An important finding of the current work is that the effects of reward magnitude upon the precision of target representation were restricted to the reaching task in which rewards were available. The lack of enhanced spatial representation in the vision-only task suggests either that sensorimotor processing or the reward context are crucial drivers of sharpened neural representation. Previous work on a perceptual discrimination task by Tankelevitch et al. (2020) provides indirect evidence in favour of a sensorimotor explanation, since these authors found an enhanced neural representation of stimulus location that persists beyond the behavioural relevance of the stimulus-reward association. In that study, participants were trained to discriminate high- and low-reward visual stimuli from non-rewarding stimuli on the basis of shape and colour features. Once participants had learned the stimulus-reward associations, they completed an orientation discrimination task involving the (now-irrelevant) reward stimuli. Crucially, the visual location of high-reward stimuli was represented more precisely (via a multivariate MEG decoding approach) than those of low-reward stimuli, both in the training and orientation-discrimination blocks. Performance on the visual discrimination task was also better when discrimination targets were presented at the same location as previously rewarded stimuli. These results suggest that, once established, the facilitation of the spatial representation of rewarding stimuli can endure after rewards are removed. Thus, the lack of enhanced representation of targets during the visual detection task that had been previously associated with highly rewarded movements in the current study should not be due merely to the fact that there were no rewards available at the time. A plausible explanation is that the circuits in which an enhanced spatial representation is manifest are part of the process that transforms sensory information about the environment and body into motor commands. Although it is also possible that differences in task context or participant engagement contributed to the lack of generalisation in reward-driven neural enhancements, we think this is unlikely as the participants effectively performed the target detection task during these vision-only blocks.

### Does a sharpened neural representation affect motor behaviour?

The timing of enhanced spatial representation due to high prospective reward is consistent with the possibility of a causal influence on motor behaviour, as it begins well before a sensitive measure of movement initiation (∼135 ms versus ∼220ms). However, it would appear unlikely that enhanced representation contributes strongly to faster initiation of the movement, as reduced reaction times were observed in both experiments, including in the absence of an altered precision of spatial representation. This suggests that pre-target motor preparation biases might be more important determinant of movement initiation than post-target sensorimotor processing. Movement speeds were, however, faster for reaches to targets associated with larger reward, and similar for equally rewarded targets irrespective of target probability. This raises the question of whether reward-related enhancements in sensorimotor processing lead to increased movement vigour.

Interestingly, we found previously that isometric aiming movements were more vigorous towards targets that were presented more frequently in an experiment in which task success (and presumably intrinsic reward) was more likely for high frequency targets (Marinovic et al., 2017). Crucially, this vigour effect varied parametrically with the angle of the presented target from that of the most frequently presented target, but varied little as a function of the time available from target presentation to movement initiation. This suggests that people develop a spatially-distributed representation of the expected value of targets according to their reward history (see e.g. (Takikawa et al., 2002)). By contrast, movement direction biases were strongly dependent on movement preparation time. This dissociation between the effects of reward and action history on movement initiation and movement vigour is consistent with the current data. We speculate that the ventral pallidum might be implicated in post-target enhancements of reach vigour, since the firing rate of neurons in this nucleus increases upon presentation of a rewarded target, remains tonically elevated until reward delivery, and strongly influences the vigour of saccades (Tachibana and Hikosaka, 2012). Such a signal should depend more strongly on the target actually presented than on the pre-target state of preparation in sensorimotor areas, and should be well placed to boost cortical sensorimotor representations that modulate movement vigour upon presentation of targets associated with high reward.

### What are the likely neural drivers of spatial representation decoded from the scalp EEG?

The method of direction decoding used in this study involves a multivariate approach that relies on all 64 scalp electrodes, and in which independent models are generated at each discrete analysis time point. There is therefore potential for signals originating anywhere in the brain to contribute to direction decoding. The method fits variance in EEG voltages across electrodes to simulated “direction channels” that reflect canonical tuning curves for polar directions. This enforces a model structure in which the variance in EEG signals must be more similar for target locations that are close together than for targets that are distant, in line with traditional population coding conceptions of visual (e.g., (Adrian, 1941; Thompson et al., 1950; Schwartz and Simoncelli, 2001; Simoncelli and Olshausen, 2001; Fabbri et al., 2010)) and motor (e.g., (Georgopoulos et al., 1986)) representations of direction. A multivariate approach that considers electrodes from all scalp locations is motivated by observations that neuronal populations in multiple brain regions including the primary and pre-motor cortices, the cerebellum, and parietal areas are tuned to hand movement direction (Fabbri et al., 2010). However, modern conceptions of neural coding rely on the principle of computation through neural population dynamics, which emphasises that information is represented within spatiotemporal neural firing patterns (Churchland et al., 2010; Vyas et al., 2020). Our approach of independent spatial decoding for each time increment clearly lacks the capacity to capture spatiotemporal dynamics. However, the fact that decoding was more effective for more rewarded targets nonetheless implies a sharpening of the multivariate pattern of EEG activity to more precisely resemble the activity pattern that is associated with a given target direction.

### What are the implications of the conventional ERP results?

There were no effects of value manipulations on with P1 or N1 ERP components contralateral to the side of target presentation, suggesting that classical early markers of attention orienting were insensitive to reward or action history manipulations. However, central midline ERP components were amplified by the presentation of a target associated with high reward, or the generation of a more highly rewarded movement. In particular there was increased negativity at the Cz electrode at timings consistent with target processing and movement preparation, and an increased positivity at the Pz electrode at mid-late movement execution. Neither of these effects were observed for movements to targets with higher probability of presentation. The Cz electrode component effect matches approximately with the timing of enhanced sharpened directional representation identified by the decoding approach, although the functional correspondence between a change in amplitude at specific electrodes and a multivariate pattern is not clear. The late parietal electrode component was dramatically suppressed during the vision only task, suggesting that sensorimotor processing or an action contingency is important. This Pz component was also much later than the period of reward-enhancement identified by the multivariate decoding approach, and so would appear unlikely to be related to the precision of sensorimotor representation.

## Conclusions

The results show a clear dissociation between the neural and behavioural responses to action and reward history expectations about the value of an action. We found that an expectation of increased reward magnitude shortens initiation time, increases movement velocity, and enhances the precision of estimates of sensorimotor representation extracted from the scalp EEG. By contrast, despite shorter movement initiation time, and previous demonstration of enhanced pre-target sensorimotor preparation of actions for targets that are more frequently encountered, we found no evidence of changes in sensorimotor representation of target or movement direction for more frequent targets. Thus, early sensorimotor processing is sharpened when the magnitude of reward associated with movement to a cued target is high.

## References

Adrian E (1941) Afferent discharges to the cerebral cortex from peripheral sense organs. The Journal of physiology 100:159.

Albrecht D, Farrar S, Hamilton D (1984) Spatial contrast adaptation characteristics of neurones recorded in the cat’s visual cortex. The Journal of physiology 347:713–739.

Benjamini Y, Hochberg Y (1995) Controlling the false discovery rate: a practical and powerful approach to multiple testing. Journal of the Royal statistical society: series B (Methodological) 57:289–300.

Brouwer GJ, Heeger DJ (2009) Decoding and reconstructing color from responses in human visual cortex. Journal of Neuroscience 29:13992–14003.

Carpenter RH, Williams M (1995) Neural computation of log likelihood in control of saccadic eye movements. Nature 377:59–62.

Carroll TJ, McNamee D, Ingram JN, Wolpert DM (2019) Rapid Visuomotor Responses Reflect Value-Based Decisions. J Neurosci 39:3906–3920.

Chen J, Scotti PS, Dowd EW, Golomb JD (2021) Neural representations of task-relevant and task-irrelevant features of attended objects. bioRxiv:2021.2005. 2021.445168.

Churchland MM, Cunningham JP, Kaufman MT, Ryu SI, Shenoy KV (2010) Cortical preparatory activity: representation of movement or first cog in a dynamical machine? Neuron 68:387–400.

Daw ND, O’Doherty JP, Dayan P, Seymour B, Dolan RJ (2006) Cortical substrates for exploratory decisions in humans. Nature 441:876–879.

Di Russo F, Martínez A, Hillyard SA (2003) Source analysis of event-related cortical activity during visuo-spatial attention. Cerebral cortex 13:486–499.

Fabbri S, Caramazza A, Lingnau A (2010) Tuning curves for movement direction in the human visuomotor system. Journal of Neuroscience 30:13488–13498.

Foster JJ, Thyer W, Wennberg JW, Awh E (2021) Covert attention increases the gain of stimulus-evoked population codes. Journal of Neuroscience 41:1802–1815.

Galaro JK, Celnik P, Chib VS (2019) Motor cortex excitability reflects the subjective value of reward and mediates its effects on incentive-motivated performance. Journal of Neuroscience 39:1236–1248.

Georgopoulos AP, Schwartz AB, Kettner RE (1986) Neuronal population coding of movement direction. Science 233:1416–1419.

Glazer JE, Kelley NJ, Pornpattananangkul N, Mittal VA, Nusslock R (2018) Beyond the FRN: Broadening the time-course of EEG and ERP components implicated in reward processing. International Journal of Psychophysiology 132:184–202.

Hickey C, Chelazzi L, Theeuwes J (2010) Reward changes salience in human vision via the anterior cingulate. Journal of Neuroscience 30:11096–11103.

Hillyard SA, Anllo-Vento L (1998) Event-related brain potentials in the study of visual selective attention. Proceedings of the National Academy of Sciences 95:781–787.

Howard IS, Ingram JN, Wolpert DM (2009a) A modular planar robotic manipulandum with end-point torque control. J Neurosci Methods 181:199–211.

Howard IS, Ingram JN, Wolpert DM (2009b) A modular planar robotic manipulandum with end-point torque control. J Neurosci Methods 181:199–211.

Jasper HH (1958) The ten twenty electrode system of the International Federation. Electroenceph Clin Neurophysiol 10:371–375.

Kok P, Jehee JF, De Lange FP (2012) Less is more: expectation sharpens representations in the primary visual cortex. Neuron 75:265–270.

Liston DB, Stone LS (2008) Effects of prior information and reward on oculomotor and perceptual choices. Journal of Neuroscience 28:13866–13875.

Maffei L, Fiorentini A, Bisti S (1973) Neural correlate of perceptual adaptation to gratings. Science 182:1036–1038.

Manohar SG, Chong TT, Apps MA, Batla A, Stamelou M, Jarman PR, Bhatia KP, Husain M (2015) Reward Pays the Cost of Noise Reduction in Motor and Cognitive Control. Curr Biol 25:1707–1716.

Marinovic W, Poh E, de Rugy A, Carroll TJ (2017) Action history influences subsequent movement via two distinct processes. Elife 6.

Maris E, Oostenveld R (2007) Nonparametric statistical testing of EEG-and MEG-data. Journal of neuroscience methods 164:177–190.

Mawase F, Lopez D, Celnik PA, Haith AM (2018) Movement repetition facilitates response preparation. Cell reports 24:801–808.

Meadows CC, Gable PA, Lohse KR, Miller MW (2016) The effects of reward magnitude on reward processing: An averaged and single trial event-related potential study. Biological Psychology 118:154–160.

Milstein DM, Dorris MC (2007) The Influence of Expected Value on Saccadic Preparation. Journal of Neuroscience 27:4810–4818.

Movshon JA, Lennie P (1979) Pattern-selective adaptation in visual cortical neurones. Nature 278:850–852.

Niv Y, Edlund JA, Dayan P, O’Doherty JP (2012) Neural prediction errors reveal a risk-sensitive reinforcement-learning process in the human brain. Journal of Neuroscience 32:551–562.

Noorbaloochi S, Sharon D, McClelland JL (2015) Payoff information biases a fast guess process in perceptual decision making under deadline pressure: evidence from behavior, evoked potentials, and quantitative model comparison. Journal of Neuroscience 35:10989–11011.

Pleger B, Blankenburg F, Ruff CC, Driver J, Dolan RJ (2008) Reward facilitates tactile judgments and modulates hemodynamic responses in human primary somatosensory cortex. Journal of Neuroscience 28:8161–8168.

Reuter E-M, Pearcey GE, Carroll TJ (2018a) Greater neural responses to trajectory errors are associated with superior force field adaptation in older adults. Experimental gerontology 110:105–117.

Reuter EM, Marinovic W, Beikoff J, Carroll TJ (2018b) It Pays to Prepare: Human Motor Preparation Depends on the Relative Value of Potential Response Options. Neuroscience 374:223–235.

Scheidt RA, Reinkensmeyer DJ, Conditt MA, Rymer WZ, Mussa-Ivaldi FA (2000) Persistence of motor adaptation during constrained, multi-joint, arm movements. J Neurophysiol 84:853–862.

Schwartz O, Simoncelli EP (2001) Natural signal statistics and sensory gain control. Nature neuroscience 4:819–825.

Serences JT, Saproo S (2010) Population response profiles in early visual cortex are biased in favor of more valuable stimuli. Journal of neurophysiology 104:76–87.

Simoncelli EP, Olshausen BA (2001) Natural image statistics and neural representation. Annual review of neuroscience 24:1193–1216.

Squires NK, Squires KC, Hillyard SA (1975) Two varieties of long-latency positive waves evoked by unpredictable auditory stimuli in man. Electroencephalography and clinical neurophysiology 38:387–401.

Storey JD (2002) A direct approach to false discovery rates. Journal of the Royal Statistical Society Series B: Statistical Methodology 64:479–498.

Summerside EM, Shadmehr R, Ahmed AA (2018) Vigor of reaching movements: reward discounts the cost of effort. Journal of neurophysiology 119:2347–2357.

Tachibana Y, Hikosaka O (2012) The primate ventral pallidum encodes expected reward value and regulates motor action. Neuron 76:826–837.

Takikawa Y, Kawagoe R, Itoh H, Nakahara H, Hikosaka O (2002) Modulation of saccadic eye movements by predicted reward outcome. Exp Brain Res 142:284–291.

Tang MF, Smout CA, Arabzadeh E, Mattingley JB (2018) Prediction error and repetition suppression have distinct effects on neural representations of visual information. Elife 7:e33123.

Tankelevitch L, Spaak E, Rushworth MF, Stokes MG (2020) Previously reward-associated stimuli capture spatial attention in the absence of changes in the corresponding sensory representations as measured with MEG. Journal of Neuroscience 40:5033–5050.

Teasdale N, Bard C, Fleury M, Young DE, Proteau L (1993) Determining movement onsets from temporal series. Journal of motor behavior 25:97–106.

Thompson J, Woolsey C, Talbot S (1950) Visual areas I and II of cerebral cortex of rabbit. Journal of Neurophysiology 13:277–288.

Tsay JS, Kim HE, Saxena A, Parvin DE, Verstynen T, Ivry RB (2022) Dissociable use-dependent processes for volitional goal-directed reaching. Proceedings of the Royal Society B 289:20220415.

Vyas S, Golub MD, Sussillo D, Shenoy KV (2020) Computation through neural population dynamics. Annual review of neuroscience 43:249–275.

